# Species-specific mitochondria dynamics and metabolism regulate the timing of neuronal development

**DOI:** 10.1101/2021.12.27.474246

**Authors:** Ryohei Iwata, Pierre Casimir, Emir Erkol, Leïla Boubakar, Mélanie Planque, Martyna Ditkowska, Katlijn Vints, Suresh Poovathingal, Vaiva Gaspariunaite, Matthew Bird, Nikky Corthout, Pieter Vermeersch, Kristofer Davie, Natalia V. Gounko, Stein Aerts, Bart Ghesquière, Sarah-Maria Fendt, Pierre Vanderhaeghen

## Abstract

The evolution of species involves changes in the timeline of key developmental programs. Among these, neuronal development is considerably prolonged in the human cerebral cortex compared with other mammals, leading to brain neoteny. Here we explore whether mitochondria influence the species-specific properties of cortical neuron maturation. By comparing human and mouse cortical neuronal maturation at high temporal and cell resolution, we found a slower pattern of mitochondria development in human cortical neurons compared with the mouse, together with lower mitochondria metabolic activity, particularly oxidative phosphorylation. Stimulation of mitochondria metabolism in human neurons resulted in accelerated maturation, leading to excitable and complex cells weeks ahead of time. Our data identify mitochondria as important regulators of the pace of neuronal development underlying human-specific features of brain evolution.

## Introduction

Developmental processes display species-specific differences in timeline, or heterochrony, which can lead to extensive divergence in organ size, cell composition or function ^1,2^. A striking illustration thereof can be found during human brain development, which is characterized by a prolonged timing of maturation of cortical neurons compared to other species ^3^. The resulting neoteny may constitute a key evolutionary mechanism enabling the exceptional gain of complexity of human brains. Human and primate cortical neurons derived from pluripotent stem cells (PSC) and xenotransplanted in the mouse cortex develop along their species-specific timeline ^4–6^, pointing to cell-intrinsic mechanisms as major players in the timing of neuronal development. The underlying mechanisms remain mostly unknown, except for human-specific genes that influence the timing of cortical neuron synaptogenesis ^7^. On the other hand, metabolism and mitochondria have emerged as drivers of cell fate transitions in many systems ^8–11^, including neural development during which mitochondria dynamics is causally linked to neurogenesis ^12,13^. Species differences in mitochondria and related metabolism have been related to lifespan, but not to developmental timing ^14,15^. Conversely, mitochondria have been linked to neuronal development and plasticity ^16–19^, but not in the context of evolution.

Could metabolism regulate the timing of cortical neuronal development, and if so in a species-specific way? To test this we first compared mitochondria dynamics and function in two species, mouse and human, whch display a very different timeline of cortical neuron maturation.

### Mitochondria growth and dynamics during cortical neuronal maturation follow a species-specific timeline

A major challenge to study neuronal maturation is that neurogenesis is not synchronous, so that populations of neurons born at different time-points co-exist at the same stage of brain development. To study cortical neuronal development with optimal temporal and cellular resolution, we developed a genetic neuronal birthdating system (NeuroD1-dependent Newborn Neuron (NNN) labeling), combining the expression of inducible CreERT2 under the control of NeuroD1 promoter, together with Cre-dependent (floxed) reporters eGFP and/or truncated CD8 (tCD8), enabling identification or magnetic-activated cell sorting (MACS) purification of the labeled cells (Figure 1A). As the NeuroD1 promoter is turned on transiently at the time of neuron generation, a pulse of Tamoxifen (4-OHT) allows the selective labeling of a cohort of neurons born at precise time-points, as assessed by EdU nuclear labeling (Figure S1A), which display a timely progression in dendritic growth and complexity (Figure 1B). To maximize the number of neuronal cells born at similar timepoints, the cultures are treated simultaneously with the Gamma-secretase inhibitor DAPT (Figure 1C), which inhibits Notch signalling and thereby increases cortical neurogenesis, followed by CD8^+^ MACS purification. The purified NNN-labeled neurons thus display homogeneous maturation patterns as assessed by scRNA-seq analyses (Figure 1C, Figure S1B).

**Fig. 1.**
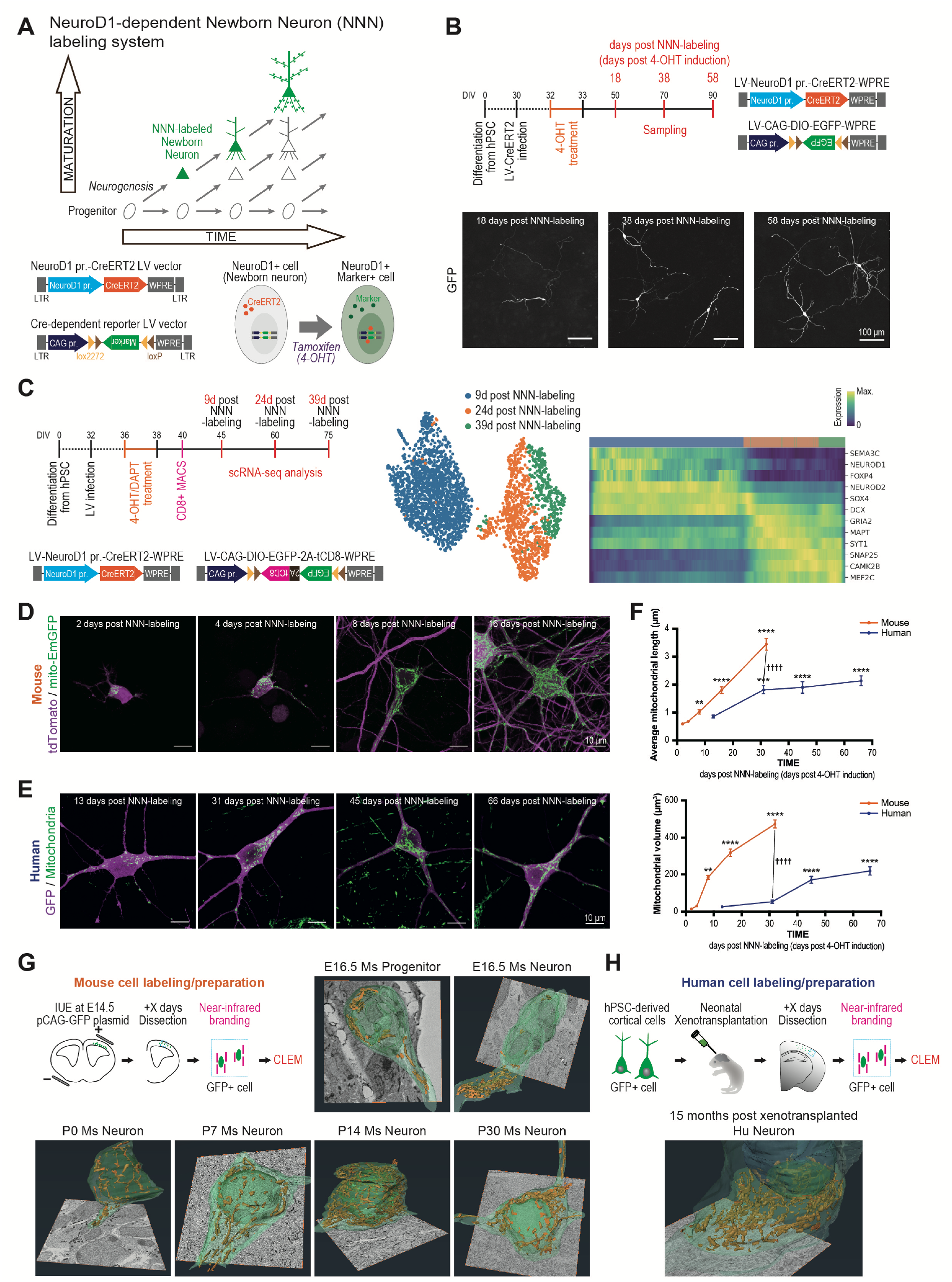
Mitochondria growth and dynamics follow a species-specific timeline during cortical neuronal maturation. (A) Schematic of NeuroD1-dependent Newborn Neuron (NNN)-labeling system. (B) Representative images following NNN-labeling in human PSC-derived cortical neurons. (C) Left: timeline of NNN-labeling for scRNA-seq analysis. Middle: UMAP of MACS-purified NNN-labeled cells annotated by their time points. Right: Heatmap of neuronal maturation related gene expression after Palantir pseudo-time analysis. Cells are ordered based on their pseudo-time values along X axis. (D,E) Representative images of mitochondrial morphology in NNN-labeled neurons in (D) mouse and (E) human. (F) Quantification of average mitochondrial length (top) and mitochondria volume (bottom) per cell from 2-3 biological replicate experiments. Data are shown as mean ± SEM. Mouse (2, 4, 8, 16, 32 days post NNN-labeling. Length: N=10, 11, 10, 10, 5 cells. Volume: N=6, 8, 10, 9, 5 cells). Human (13, 31, 45, 66 days post NNN-labeling. Length: N=20, 15, 17, 18 cells. Volume: N=9, 11, 22, 20 cells). Dunnett’s multiple comparisons test. Compared with first time point: ***P < 0.001, ****P < 0.0001. Compared mouse and human: Unpaired t test. ††††P < 0.0001. (G) Schematic of mouse (Ms) cortical neuron birth-dating by in utero electroporation (IUE) followed by correlative light and electron microscopy (CLEM), and representative images at indicated time-point following IUE. E: embryonic day. P: postnatal day. Yellow structure: Mitochondria. Bottom image: 20μm square. (H) Schematic of human (Hu) cortical neuron birth-dating followed by neonatal mouse cortex xenotransplantation used for CLEM, and representative image at 15 months post-transplantation. Only a small part of the human neuron is modeled as a representative piece. Bottom image: 20μm square.

By applying the NNN-labeling system to mouse cortical neurons in combination with mitochondria labeling with Mito-EmGFP (Emerald-GFP protein fused to mitochondrial targeting sequence of COX8A) (Figure 1D), we observed that mitochondria are initially small in size and relatively sparse in newly born mouse neurons, as expected ^12^, and gradually grow in 2-3 weeks during neuronal maturation (Figure 1D,F). Surprisingly however, similar examination of PSC-derived human cortical neurons revealed a much more prolonged timeline of mitochondria development spanning several months (Figure 1E,F).

We next looked at mitochondria dynamics in mouse and human developing cortical neurons in vivo. We first labeled cortical neurons with eGFP using in utero electroporation (IUE) at E14.5, followed by correlative light and electron microscopy (CLEM) analysis at several time points until cortex maturation at postnatal day 30 (P30) (Figure 1G). This confirmed that mitochondria gradually reach maximal levels of growth and size in 2-3 weeks in mouse cortical neurons in vivo (Figure 1G). On the other hand, we performed xenotransplantation of DAPT-treated PSC-derived human cortical neurons transduced with eGFP. In this system, xenotransplanted neurons integrate in mouse cortex within a few days after transplantation and display a protracted pattern of maturation, taking >12 months to reach maturation ^5^. CLEM performed on these neurons revealed that the human cortical neurons displayed much slower patterns of mitochondria growth than their mouse counterparts, taking >12 months to reach similar levels of mitochondria development (Figure 1H).

These data indicate that mitochondria morphological development follows a species-specific timeline that is highly correlated with neuronal maturation.

### Mitochondria metabolic activity is lower in human than mouse developing neurons

Next, we examined the functional properties of mitochondria during neuronal development, first focusing on mitochondrial oxidative phosphorylation (OXPHOS) and electron transport chain (ETC) capacity. We measured mitochondrial oxygen consumption rate (OCR) using Oxygraphy ^20^ on highly enriched preparations (>95% neurons) of birth-timed mouse and human cortical neurons (Figure 2A; Figure S2A). We found that mitochondrial OCR in human and mouse cortical neurons gradually increases along species-specific timelines, in line with their morphological development (Figure 2B-D, Figure S2B-D). Most strikingly, stimulated OCR was found to be >10 times higher in mouse than human neurons (Figure 2C,D, Figure S2), indicating much higher levels of ETC in mouse than in human neurons. We next performed mass spectrometry-based ^13^C tracer analysis ^21^ of enriched preparations (>95% neurons) of human and mouse cortical neurons (>95% neurons, Figure S3A) at a similar age (9-12 days following neuronal birth), focusing on glucose metabolism (Figure 3). On the one hand, this revealed a higher enrichment of lactate from ^13^C_6_-glucose and a higher secretion of lactate in human compared with mouse neurons (Figure 3A-C, Figure S3B,C). On the other hand, the mitochondria-driven use of pyruvate through the tricarboxylic acid (TCA) cycle was comparatively lower than in mouse neurons (Figure 3B). Collectively these data indicate that human developing cortical neurons display lower levels of mitochondria-driven TCA and oxidative activity than their mouse counterparts at similar birth-time.

**Fig. 2.**
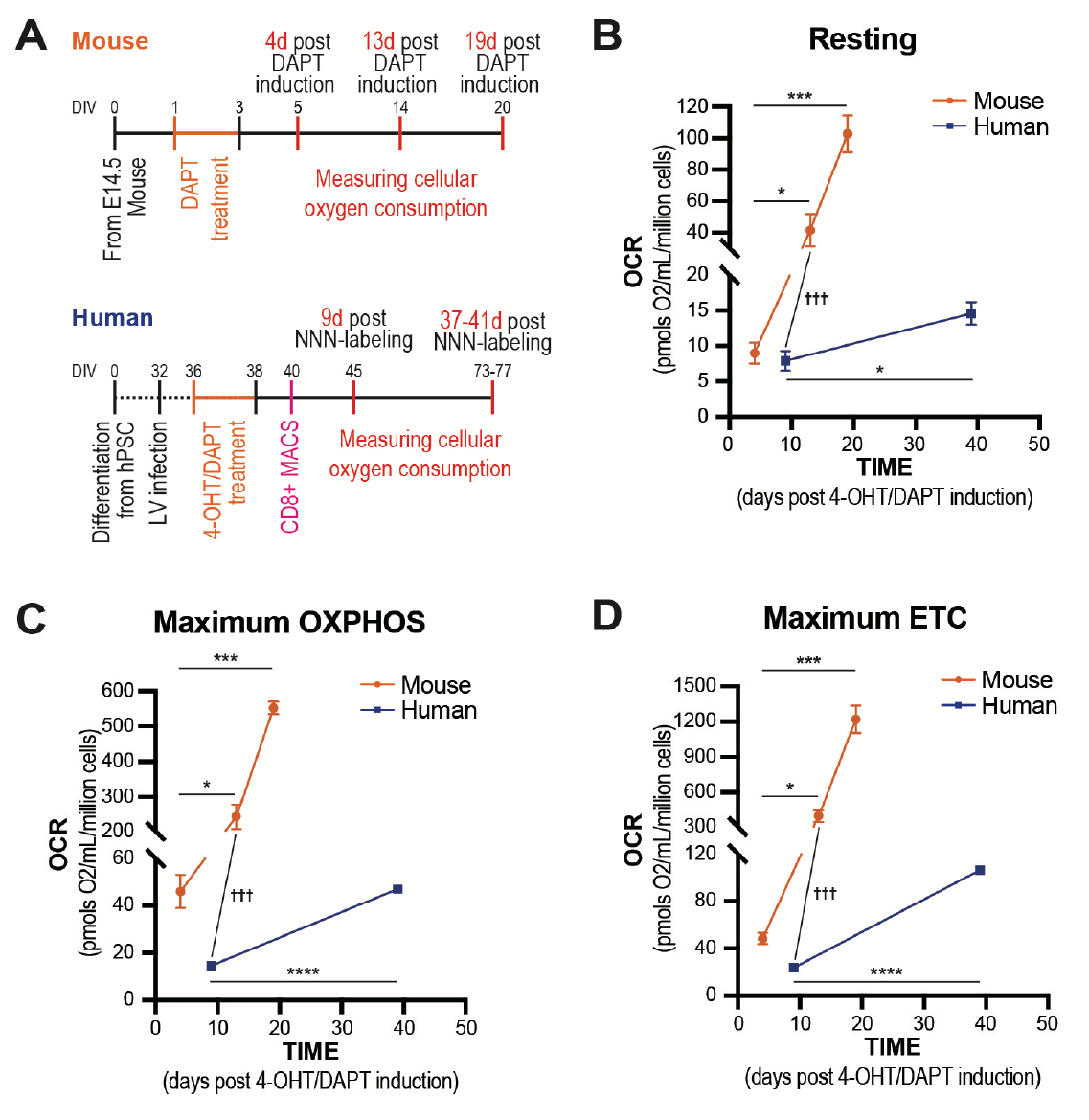
Mitochondria OXPHOS activity increases during neuronal maturation and displays human-mouse interspecies differences. (A) Timeline of experiments to measure mitochondria oxidative activity in mouse and human neurons. Human neurons were further purified from progenitors using MACS, (B-D) Quantification of oxygen consumption rate (OCR) during neuronal development. Data are shown as mean ± SEM from at least two biological replicates. (B) Resting OCR. (C) Maximum oxidative phosphorylation (OXPHOS) capacity under coupled condition. (D) Maximum electron transport chain (ETC) capacity under uncoupled condition (uncoupler CCCP treated). Mouse (4, 13, 19 days post NNN-labeling. N=6, 8, 4). Human (9, 37-41 days post NNN-labeling. N=6, 7). Mouse: Dunn’s multiple comparisons test. Human: Unpaired t test. Mouse vs Human: Mann Whitney test. *P < 0.05, **P < 0.01, *** or †††P < 0.001, ****P < 0.0001.

**Fig. 3.**
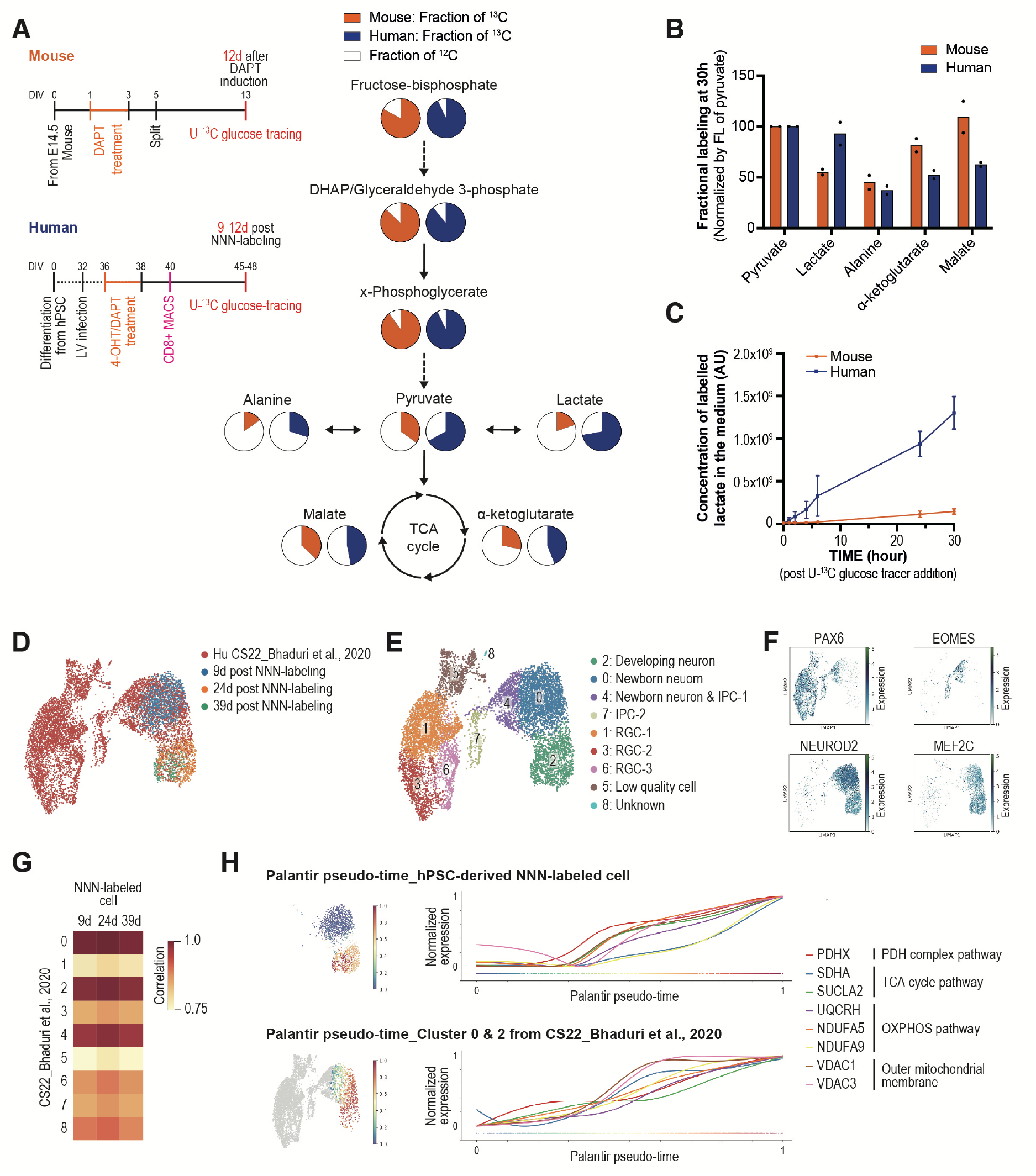
Species-specific metabolism features of cortical neurons. (A) Left: timeline of experiments to measure glycolysis and TCA cycle intermediate metabolites labeling by U-^13^C-labeled glucose at 30h post tracer addition. Human neurons were further purified from progenitors using MACS. Right: ^13^C enrichment patterns of the metabolites in (Orange) Mouse and (Blue) Human. DHAP: Dihydroxyacetone phosphate. (B) Normalized fractional labeling of metabolites by labelled fraction of pyruvate at 30h post tracer addition. Each data point represents an independent biological replicate. FL: fractional labeling. (C) Concentration of ^13^C-labelled lactate in the culture medium at different time points. Data are shown as mean ± SEM. Mouse: N=2, Human: N=5. AU: arbitrary units. (D-F) UMAP of NNN-labeled human cortical neurons in vitro, and in vivo human cortical cells from fetal stage Carnegie (CS) 22 fetal stage ^22^ after Harmony batch correction. (E) Unsupervised Leiden clustering after integration with Harmony algorithm. (F) Normalized expressions of PAX6, EOMES, NEUROD2 and MEF2C were shown as radial glial cells (RGC), intermediate progenitor cells (IPC), excitatory neurons and developing neuron markers, respectively. (G) Heatmap of Pearson correlation coefficients per time point of the mean pathway scores of metabolic pathways (Glycolysis, PDH complex, TCA, OXPHOS, Table. S1) from NNN-labeled neurons and CS22 in vivo cortex clusters identified by Leiden algorithm. (H) Expression trends of representative metabolic pathway genes are plotted along the Palantir pseudo-time axis for NNN-labeled neurons (Top) and CS22 neurons from cluster 0 and 2 (Bottom).

Our oxygraphy data point to a striking temporal patterning of mitochondria metabolism during neuronal maturation. To gain insights on the upstream mechanisms and in vivo relevance of these observations, we examined our scRNA-seq of in vitro birth-dated human neurons, together with in vivo human fetal cortex scRNAseq datasets ^22^ (Figure 3D-F, Figure S4), focusing on mitochondria and metabolism genes (Figure 3G,H). This revealed a temporal increase of mitochondria metabolism-related genes including TCA cycle and OXPHOS pathways during neuronal maturation (Figure 3H), thus highly consistent with the oxygraphy data. Importantly, the in vitro and in vivo patterns of expression of mitochondria/metabolic genes were highly similar and correlated (Figure 3G,H), thus providing direct in vivo validation for our in vitro observations.

### Increasing mitochondria activity in human neurons leads to accelerated neuronal maturation, growth and complexification

We next tested whether the species differences observed in mitochondria metabolic functions could be causally linked to the speed of neuronal maturation.

Glucose tracer experiments revealed an interspecies difference in the pyruvate to lactate conversion (Figure 3A-C), which is catalyzed by lactate dehydrogenase (LDH). LDH is a tetrameric enzyme with isoenzymes composed of different proportions of two common subunits, A and B, LDHA favoring the conversion of pyruvate to lactate, LDHB favoring the conversion of lactate to pyruvate ^23^ (Figure 4A). To boost mitochondria activity in developing neurons, we therefore targeted the activity of LDHA. As expected, treatment with a chemical inhibitor of LDHA, GSK-2837808A (GSK) ^24^, resulted in increased mitochondria oxidative activity in human neurons, as measured by mitochondria OCR (Figure 4B, Figure S5). This result is consistent with previous reports in cancer cells, in which genetic depletion of LDHA increases mitochondrial OXPHOS activity ^25^.

**Fig. 4.**
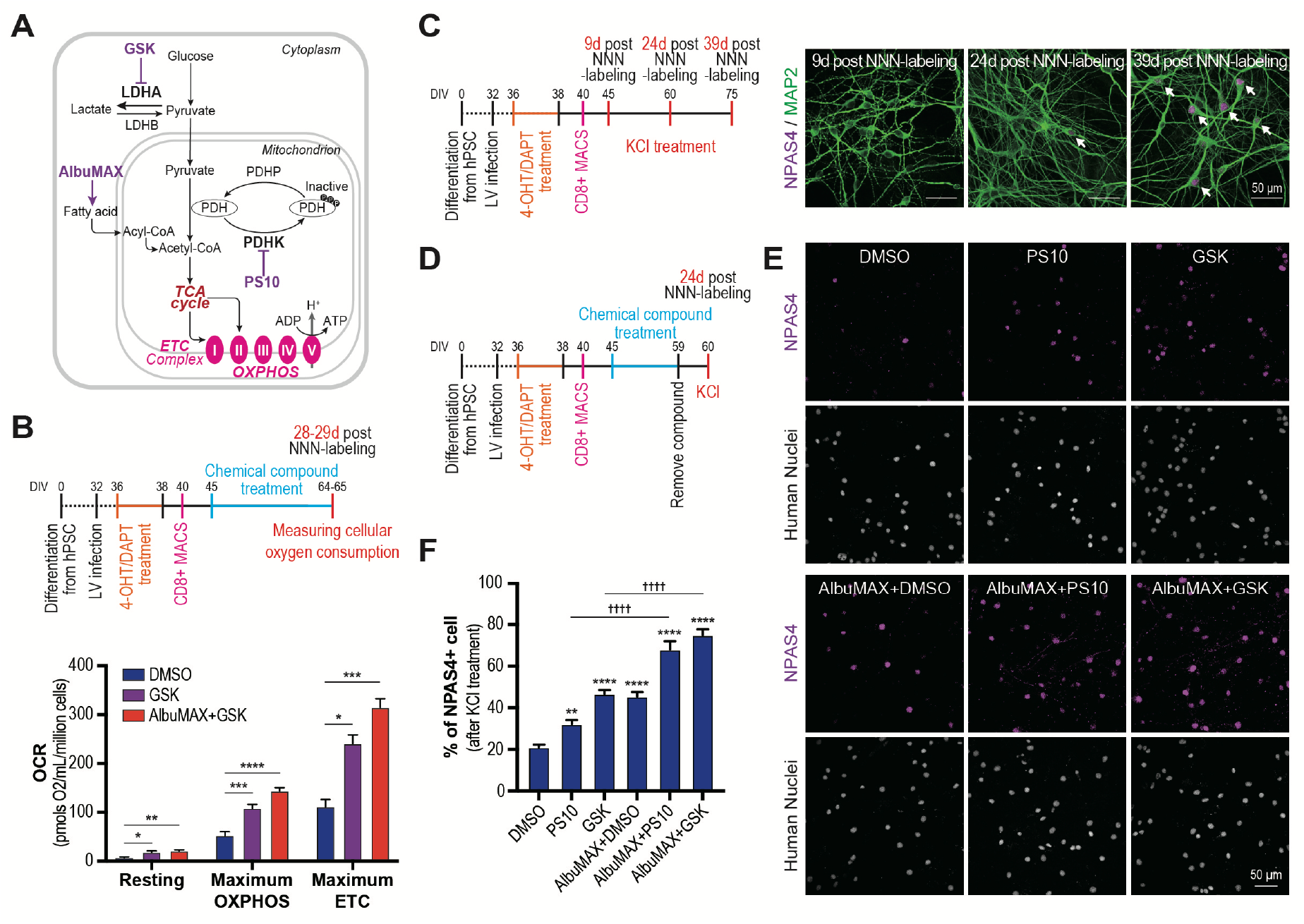
Increasing mitochondria metabolism accelerates human neuronal maturation. (A) Schematic of metabolic pathways targeted by indicated chemical compounds. GSK: GSK-2837808A. LDHA/B: lactate dehydrogenase A/B. PDHK/P: pyruvate dehydrogenase kinase/phosphatase (B) Timeline of oxygraphy experiment in human neurons following chemical treatments. Quantified OCR from two biological replicate experiments. Data are shown as mean ± SEM. Dunnett’s or Dunn’s multiple comparisons test. DMSO: N=7, GSK: N=7, AlbuMAX+GSK: N=8. (C) Experimental scheme and representative images of KCl-induced NPAS4 expression in NNN-labeled human neurons over time. Arrow: NPAS4+ neurons. (D) Experimental scheme of chemical compounds treatments. (E) Representative images of the number of NPAS4 positive neurons, among all neurons (human-specific nuclear antigen), induced by KCl following treatment with indicated chemical compounds. (F) Quantification of the proportion of NPAS4-positive neurons induced by KCl following treatment with indicated chemical compounds. Data are shown as mean ± SEM from 3 independent biological replicates. DMSO: N=20, PS10: N=20, GSK: N=15, AlbuMAX+DMSO: N=15, AlbuMAX+PS10: N120, AlbuMAX+GSK: N=20 ROIs. Tukey’s multiple comparisons test. Compare with DMSO: *P < 0.05, **P < 0.01, ***P < 0.001, **** P < 0.0001. ††††P < 0.0001.

To probe the speed of maturation we assessed the classical parameter of neuronal excitability, by measuring their ability to respond to membrane depolarization following KCl addition, using as a read-out the expression of the activity-dependent immediate early gene NPAS4 ^26^. We thus found that very few NNN-labeled human cortical neurons respond to KCl shortly after their birth (9-24 days), while most of them become responsive only weeks later (39 days) (Figure 4C).

We then examined the impact of LDHA inhibitors on human cortical neuron maturation, by treating the cells from day 9 to 23, when the neurons are still poorly responsive to KCl depolarization (Figure 4D). Importantly, the LDH inhibitor treatment was stopped for at least one day before assessing maturation: this temporal dissociation allowed to distinguish genuine effects on timing of maturation per se from an impact of mitochondria function on neuronal responsiveness to KCl. While in control conditions, few neurons at day 24 responded to KCl treatment, as expected (Figure 4E,F), prior inhibition of LDHA led to a major increase in the proportion of neurons displaying NPAS4 responses, indicating a strong acceleration of neuronal maturation in response to increased mitochondria respiration (Figure 4B, Figure S5B). Additional treatment of the neurons with preparations of free fatty acids (FFAs) (AlbuMAX), an additional fuel besides glucose for mitochondrial TCA cycle activity, led to additive increase in mitochondrial activity and neuronal maturation (Figure 4B,E,F, Figure S5), further pointing to mitochondrial metabolic activity as an important drive of the speed of neuronal maturation. A similar effect was also observed following treatment with PS10, a PDH kinase (PDHK) pan-inhibitor ^27^, a treatment that also stimulates mitochondrial metabolism by increasing the conversion of Pyruvate into Acetyl-CoA (Figure 4A,E,F). Finally, as these experiments were performed on human cortical neurons cultured on mouse astrocytes, we tested the impact of the same compounds on pure neuronal populations, which resulted in the same degree of acceleration of neuronal developmental timing (Figure S6).

Overall, these data indicate that increasing mitochondria metabolic activity results in acceleration of human neuronal differentiation.

Finally, we examined the global consequences of mitochondria-boosting treatments on neuronal size and dendritic length, as well as branching of dendritic arbors, which constitute crucial aspects of neuronal maturation. GSK/FFA-treated human cortical neurons displayed strikingly larger neuronal size and increased dendritic length and complexity (Figure 5). Remarkably, the growth and complexity pattern of 24 days-old neurons treated with GSK/FFA reached at least the same maturation level as for control neurons 58 days after their birth (Figure 1B, 5A), indicating that mitochondria boosting accelerates neuronal development towards growth and complexity several weeks ahead of time.

**Fig. 5.**
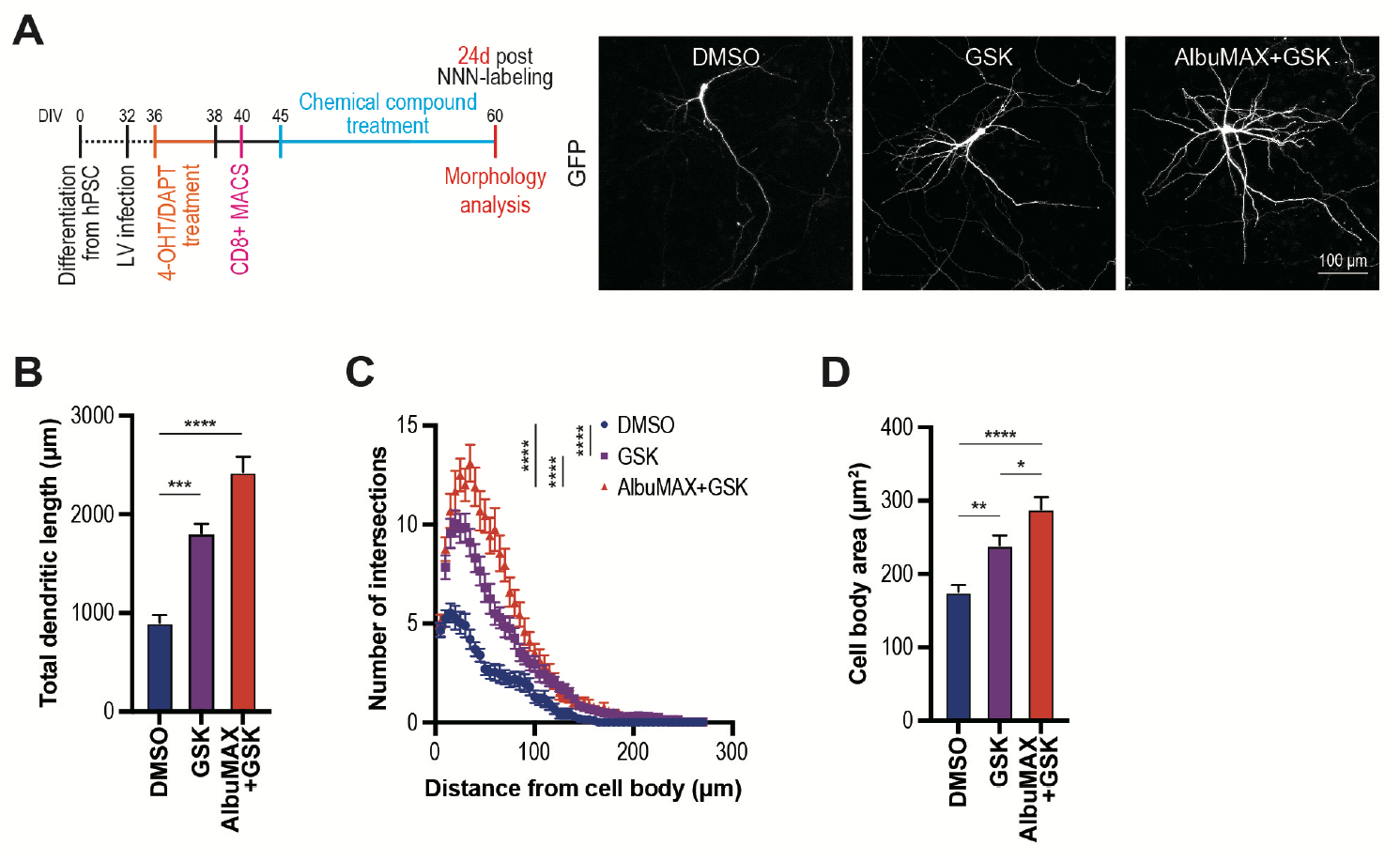
Increasing mitochondria metabolism accelerates neuronal growth and complexification. (A) Experimental scheme and representative images of NNN-labeled neurons following treatment with indicated chemical compounds. (B-D) Quantification of neuronal morphology analysis from 2 biological replicates. N=20 neurons for each condition. (B) Total dendritic length. Data are shown as mean ± SEM. Dunn’s multiple comparisons test. (C) Sholl analysis of dendritic branching. (D) Cell body area. Tukey’s multiple comparisons test. *P < 0.05, **P < 0.01, ***P < 0.001, ****P < 0.0001.

## Discussion

Altogether these data indicate that the rates of mitochondrial activity can directly influence the species-specific developmental timeline of neuronal maturation. Future work should delineate the underlying upstream mechanisms, i.e. the origin of species-differences in developmental rates of metabolism. Our transcriptomic data suggest that at least part of the temporal differences in mitochondria maturation likely originate from patterns of mitochondria gene expression, but the observed species differences in mitochondria function likely also involve post-translational mechanisms such as modifications of mitochondrial dynamics and assembly of the ETC ^28,29^.

As for downstream mechanisms, differences in mitochondrial metabolic rates could alter the species-specific turnover rate of RNAs and proteins ^30,31^, which was recently associated with differential developmental timing in other systems ^32,33^. Mitochondrial metabolism could also regulate the rate of neuronal development through the generation of specific metabolites that affect post-translational modifications of key proteins linked to chromatin remodeling, as shown in stem cell differentiation and oncogenesis ^11,34^.

Our data strongly suggest a direct link between differences in species-specific properties of cortical neurons and human brain neoteny, in line with previous reports of years-long periods of aerobic glycolysis in the developing human cortex ^35^. Aerobic glycolysis has long been thought to constitute a hallmark of cellular proliferation, as during oncogenesis ^36^. Here we find that mitochondrial OXPHOS activity regulates the speed of post-mitotic neuron morphological and functional development, and displays striking species differences. Mitochondria metabolism could thus also contribute to heterochrony in other developmental contexts and species ^1^. Importantly, while our data are focused on developmental timing, they raise the possibility that species-specific changes in mitochondria could also influence neuronal plasticity and aging, through changes in epigenetics and/or proteostasis.

Finally, our data open new avenues to speed up the tempo of human neuronal maturation in vitro in a physiological fashion using metabolic modulators. This could have important implications for the rapidly growing field of PSC-based modeling of human neurological diseases, which remains greatly hindered by the slowness of human neuron development.

## FIGURES and FIGURE LEGENDS

**Fig. S1.**
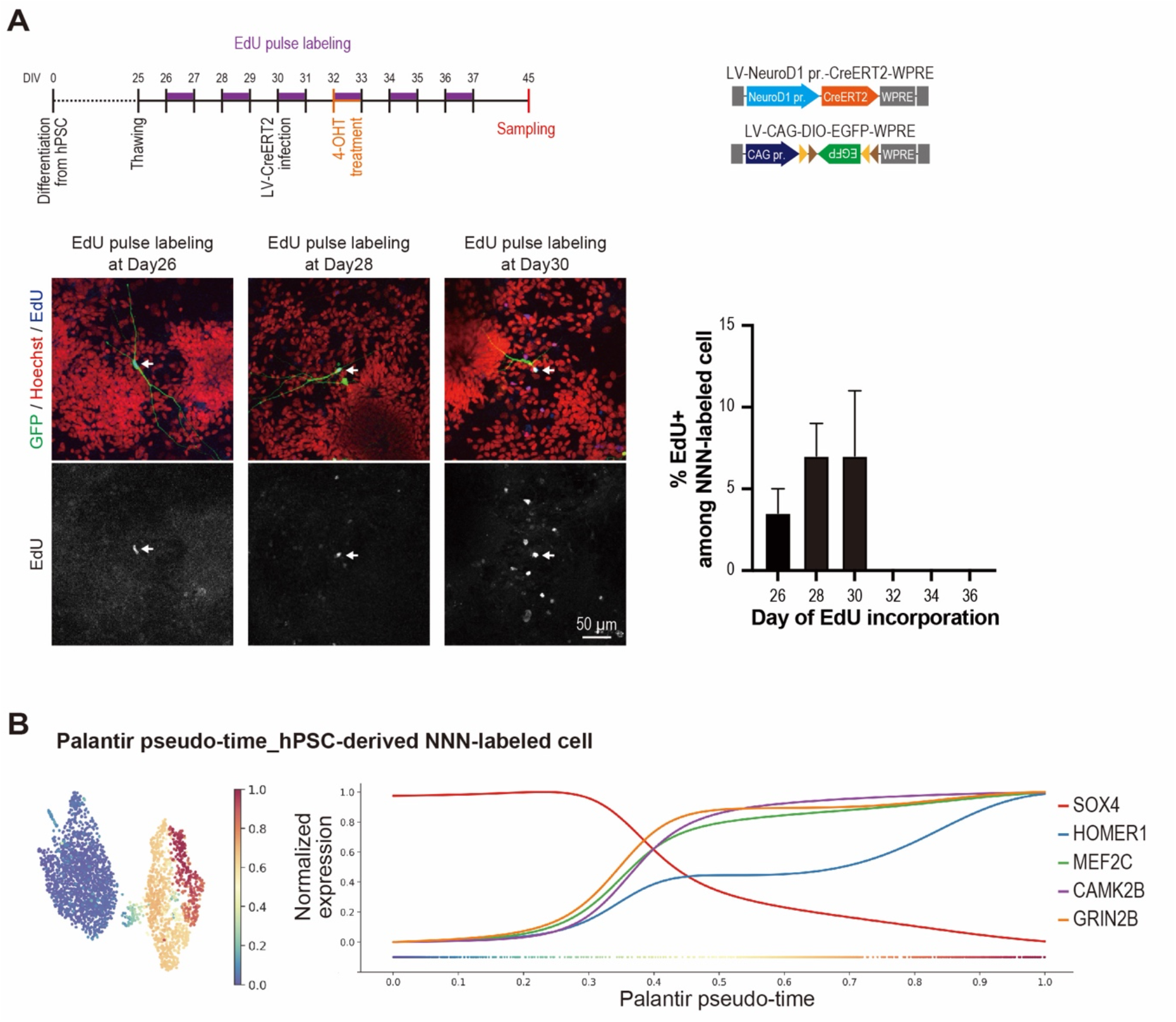
Characterization of NNN-labeling system. (A) Experimental scheme and representative images of EdU incorporated NNN-labeled neurons. Arrows: EdU+ cells. Quantification of EdU+ cell among NNN-labeled cells. N=2 biological replicates. (B) Expression trends of neuronal development marker genes are plotted along the Palantir pseudo-time axis of NNN-labeled neurons.

**Fig. S2.**
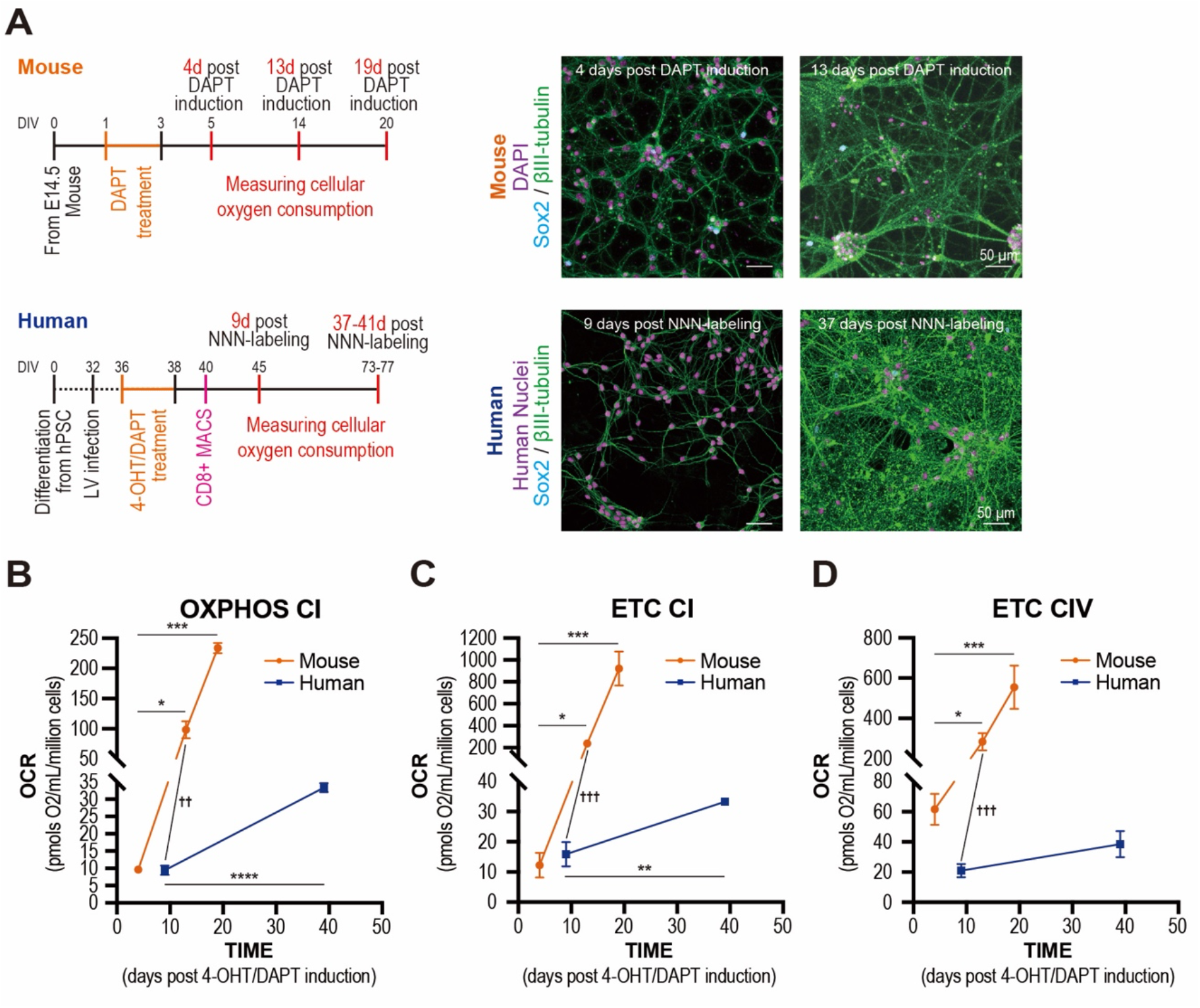
Mitochondrial OXPHOS and ETC activity in mouse and human neurons. (A) Experimental scheme and representative images of mouse and human neurons used for oxygraphy experiment. (B-D) Quantified oxygen consumption rate (OCR). Data are shown as mean ± SEM. (B) CI-linked oxidative phosphorylation (OXPHOS) capacity under coupled condition. (C) CI-linked electron transport chain (ETC) capacity under uncoupled condition (uncoupler, CCCP treated). (D) CIV-linked electron transport chain (ETC) capacity under uncoupled condition (uncoupler, CCCP treated). Mouse (4, 13, 19 days post NNN-labeling. N=6, 8, 4). Human (9, 37-41 days post NNN-labeling. OXPHOS CI: N=4, 7. ETC CI & CIV: N=6, 7). Mouse: Dunn’s multiple comparisons test. Human: Unpaired t test. Mouse vs Human: Mann Whitney test. *P < 0.05, ** or ††P < 0.01, *** or †††P < 0.001, ****P < 0.0001.

**Fig. S3.**
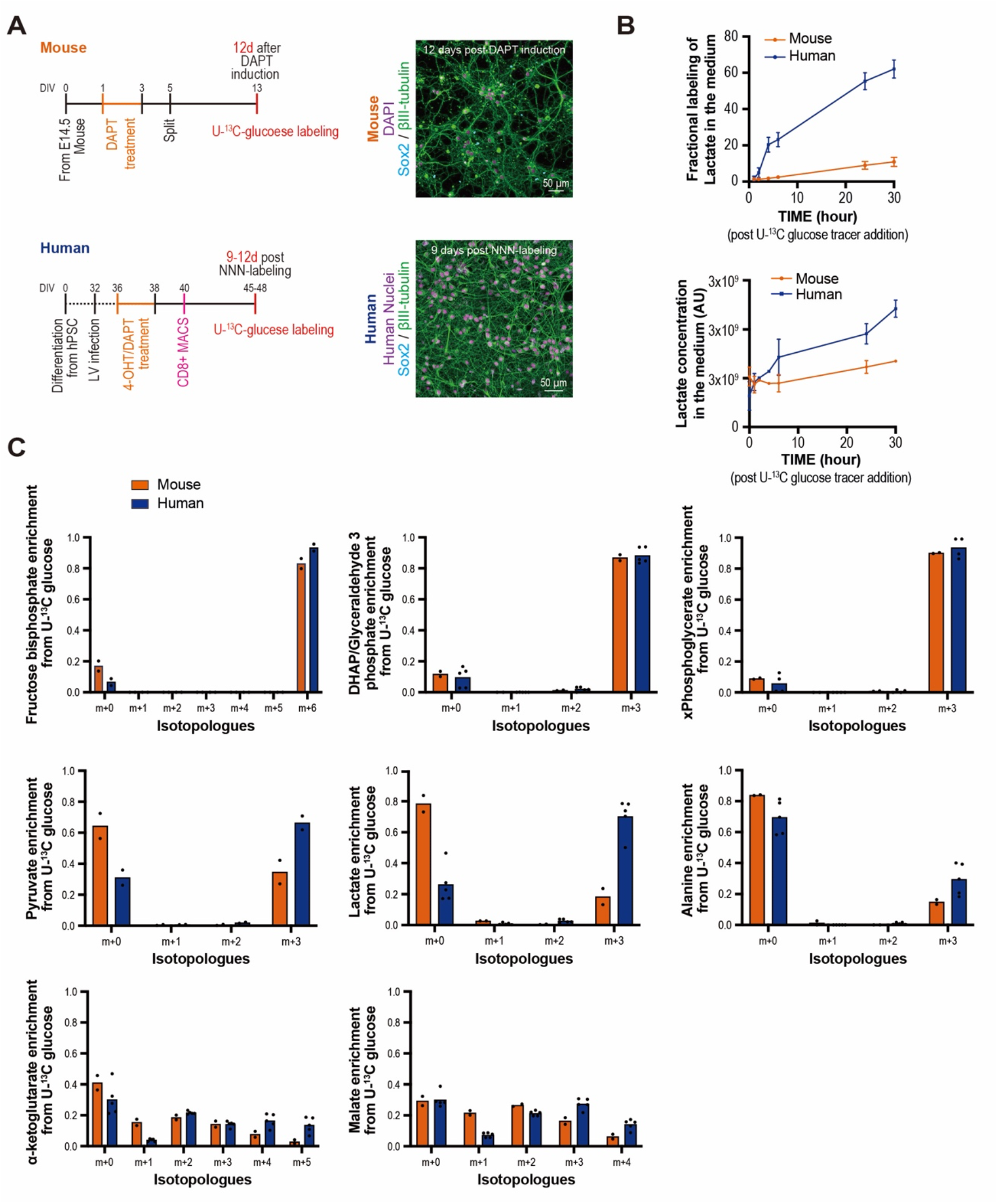
U-^13^C glucose tracing and targeted metabolites in mouse and human cortical cells. (A) Experimental scheme and representative images of mouse and human neurons that used for the ^13^C-labeled glucose-tracing experiment. (B) Top: Fractional labeling of lactate in the medium at different time points. Bottom: Lactate (^12^C and ^13^C) concentration in the medium at different time points. Data are shown as mean ± SEM. Ms: N=2, Hu: N=5. AU: arbitrary units. (C) Isotopologue distribution following ^13^C for fructose bisphosphate (Ms: N=2, Hu: N=2), glyceraldehyde 3-phosphate (Ms: N=2, Hu: N=5), x-phosphoglycerate (Ms: N=2, Hu: N=4), pyruvate (Ms: N=2, Hu: N=2), lactate (Ms: N=2, Hu: N=5), alanine (Ms: N=2, Hu: N=5), α-ketoglutarate (Ms: N=2, Hu: N=5) and malate (Ms: N=2, Hu: N=5). Each data point represents an individual sample.

**Fig. S4.**
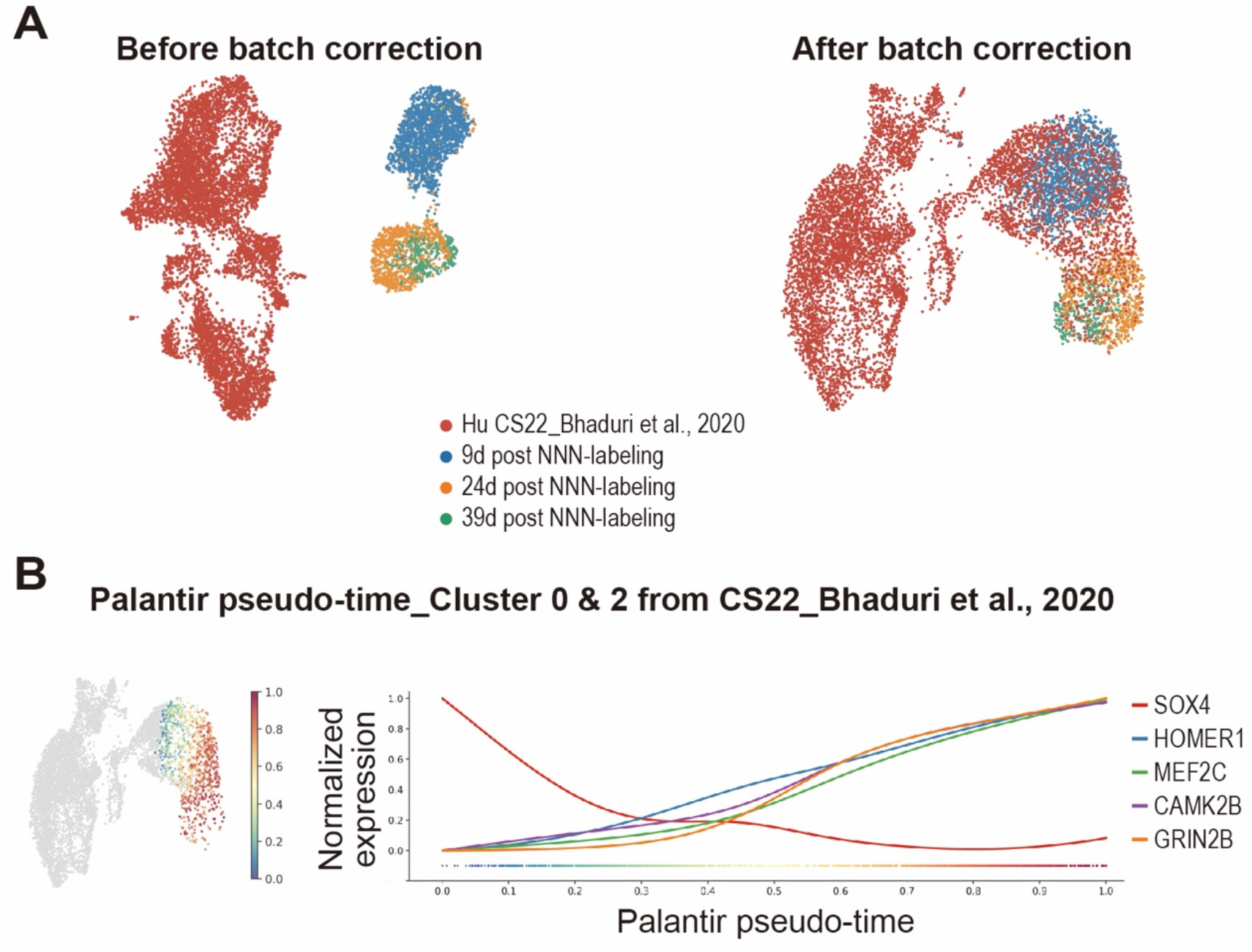
Transcriptome analysis of human in vitro and in vivo sample. (A) Left: UMAP of hPSC-derived NNN-labeled neurons and primary human cortical cells from CS22 fetal cortex before batch correction. Right: After Harmony batch correction. (B) Expression trends of neuronal development marker genes are plotted along the Palantir pseudo-time axis of cluster 0 and 2 of CS22.

**Fig. S5.**
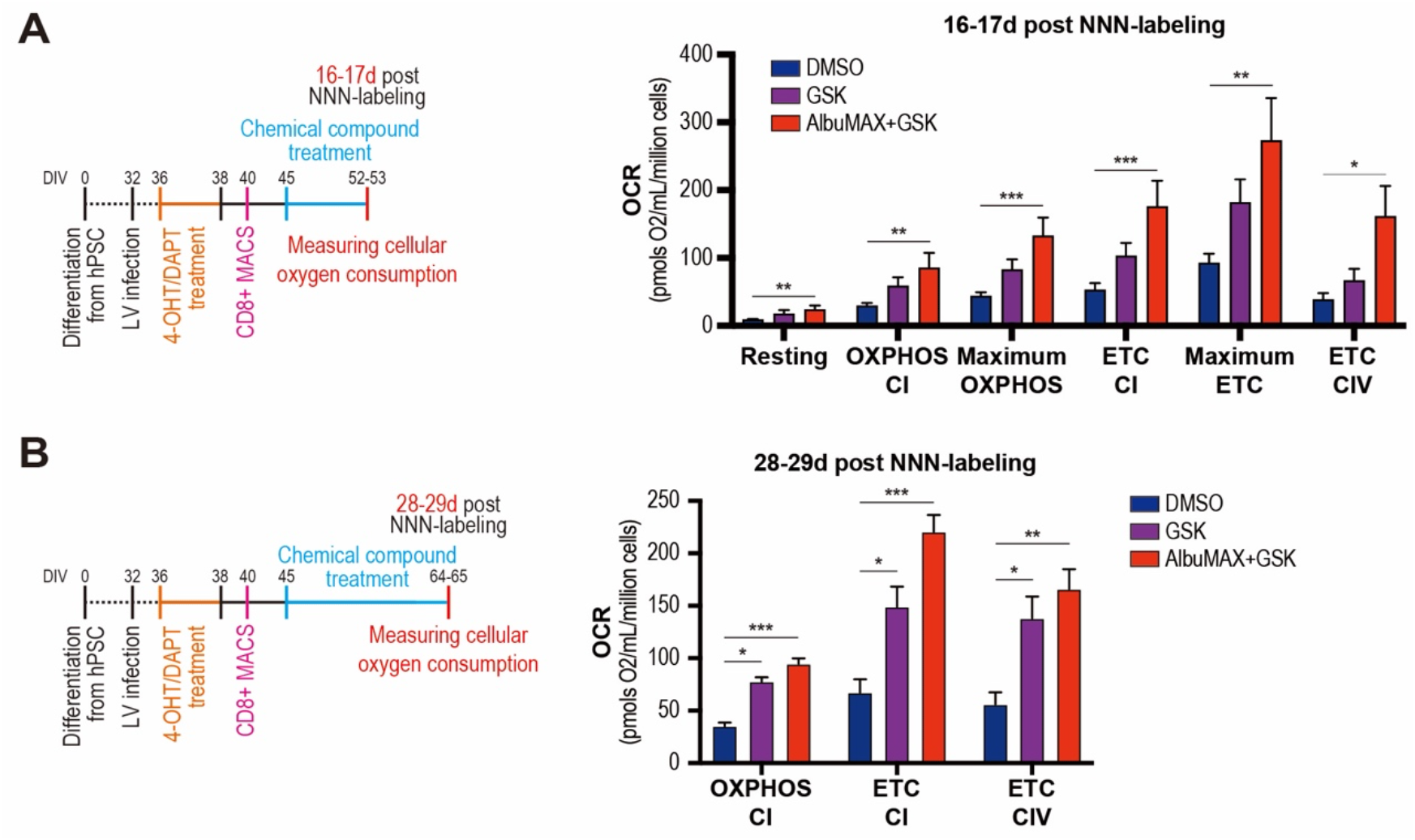
Impact of LDHA inhibition on human neuron mitochondrial activity. (A-B) Experimental scheme and quantified OCR from more than two biological replicate experiments. Data are shown as mean ± SEM. (A) 16-17 days post NNN-labeled human neurons. DMSO: N=9, GSK: N=10, AlbuMAX+GSK: N=11. (B) 28-29 days post NNN-labeled human neurons. DMSO: N=7, GSK: N=7, AlbuMAX+GSK: N=8. Dunnett’s or Dunn’s multiple comparisons test. *P < 0.05, **P < 0.01, ***P < 0.001.

**Fig. S6.**
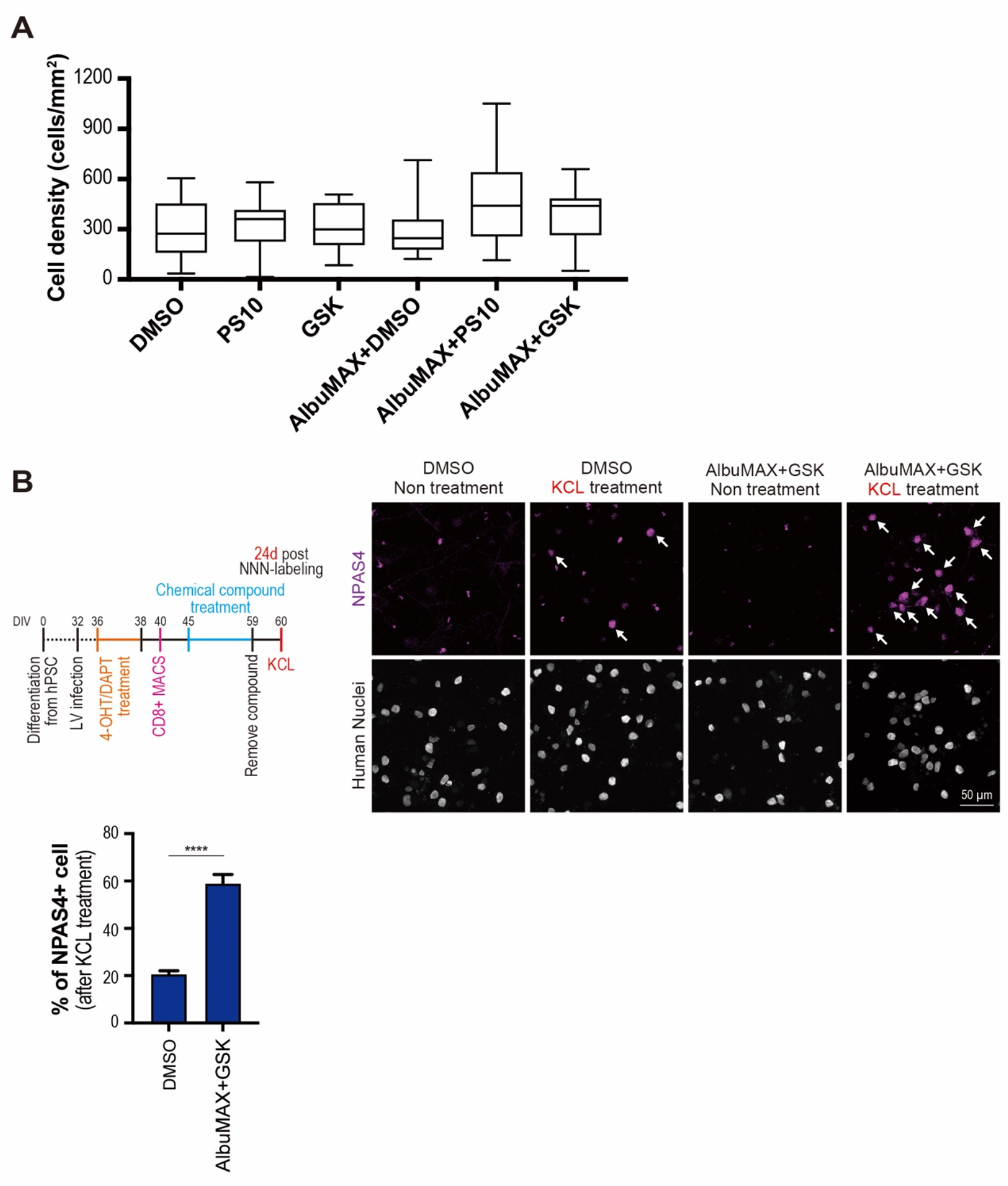
Mitochondria boosting treatment has no detectable impact on neuronal density and accelerates neuronal development in absence of astrocytes. (A) Quantified cell density and comparison among conditions. Data is represented by box plots displaying minimum to maximum values. The box extends from the 25^th^ to the 75^th^ percentile. There is no significant difference of cell density among conditions. DMSO: N=35, PS10: N=35, GSK: N=30, AlbuMAX+DMSO: N=30, AlbuMAX+PS10: N=20, AlbuMAX+GSK: N=35 ROIs. (B) Left: Experimental scheme. At day 40, MACS-sorted cells were plated on coverslip without mouse astrocytes. Right: Representative images of NNN-labeled neurons. Arrow: NPAS4+ cell. Date are shown as mean ± SEM. DMSO: N=10, AlbuMAX+GSK: N=10 ROIs. Unpaired t test. ****P < 0.0001.

**Table S1.**
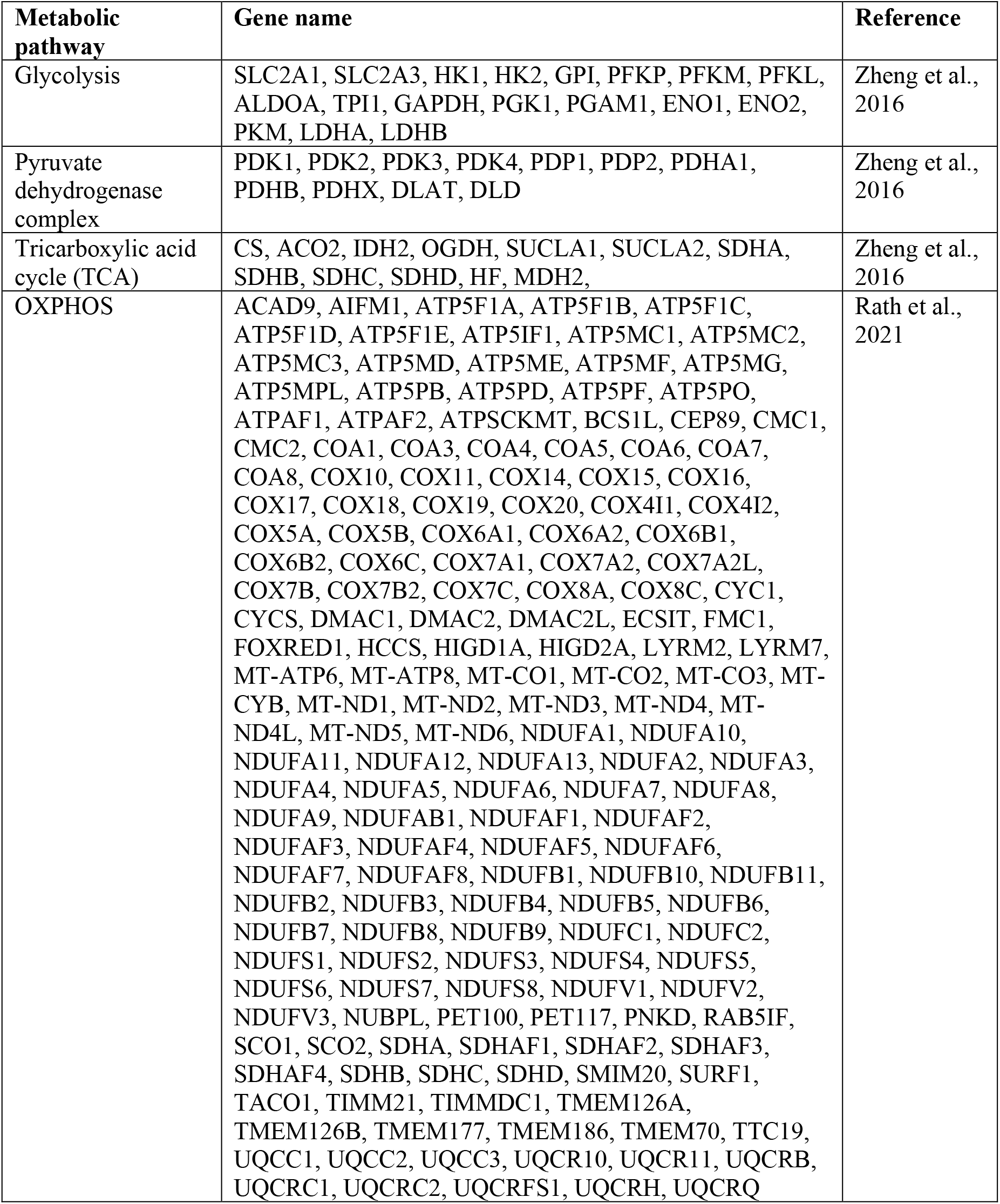
Gene list for Pearson correlation analysis.

## Acknowledgments

We thank members of the PV lab and CBD for helpful discussions and precious help.

## Author contributions

Conceptualization and Methodology, RI, PC, PV; Investigation, RI, PC, EE, LB, MD, SP, BG, SE, VG, SMF, PV.; Formal Analysis, RI, PC, EE, NC, SP, PVM, BG, KD, BG, SMF, PV; Writing, RI, PC, PV; Funding acquisition, PV; Resources, PV; Supervision, PV.

## Data and materials availability

All data are available in the manuscript or the supplementary materials. All materials are available upon request from Pierre Vanderhaeghen.

